# Exploring community structure in biological networks with random graphs

**DOI:** 10.1101/001545

**Authors:** Pratha Sah, Lisa O. Singh, Aaron Clauset, Shweta Bansal

## Abstract

**Background:** Community structure is ubiquitous in biological networks. There has been an increased interest in unraveling the community structure of biological systems as it may provide important insights into a system’s functional components and the impact of local structures on dynamics at a global scale. Choosing an appropriate community detection algorithm to identify the community structure in an empirical network can be difficult, however, as the many algorithms available are based on a variety of cost functions and are difficult to validate. Even when community structure is identified in an empirical system, disentangling the effect of community structure from other network properties such as clustering coefficient and assortativity can be a challenge.

**Results:** Here, we develop a generative model to produce undirected, simple, connected graphs with a specified degrees and pattern of communities, while maintaining a graph structure that is as random as possible. Additionally, we demonstrate two important applications of our model: (a) to generate networks that can be used to benchmark existing and new algorithms for detecting communities in biological networks; and (b) to generate null models to serve as random controls when investigating the impact of complex network features beyond the byproduct of degree and modularity in empirical biological networks.

**Conclusion:** Our model allows for the systematic study of the presence of community structure and its impact on network function and dynamics. This process is a crucial step in unraveling the functional consequences of the structural properties of biological systems and uncovering the mechanisms that drive these systems.

## Background

Network analysis and modeling is a rapidly growing area which is moving forward our understanding of biological processes. Networks are mathematical representations of the interactions among the components of a system. Nodes in a biological network usually represent biological units of interest such as genes, proteins, individuals, or species. Edges indicate interaction between nodes such as regulatory interaction, gene flow, social interactions, or infectious contacts [1]. A basic model for biological networks assumes random mixing between nodes of the network. The network patterns in real biological populations, however, are typically more heterogeneous than assumed by these simple models [2]. For instance, biological networks often exhibit properties such as degree heterogeneity, assortative mixing, non-trivial clustering coefficients, and community structure (see review by Proulx et al. [1]). Of particular interest is community structure, which reflects the presence of large groups of nodes that are typically highly connected internally but only loosely connected to other groups [3,4]. This pattern of large and relatively dense subgraphs is called assortative community structure. In empirical networks, these groups, also called modules or communities, often correspond well with experimentally-known functional clusters within the overall system. Thus, community detection, by examining the patterns of interactions among the parts of a biological system, can help identify functional groups automatically, without prior knowledge of the system’s processes.

Although community structure is believed to be a central organizational pattern in biological networks such as metabolic [5], protein [6,7], genetic [8], food-web [9,10] and pollination networks [11], a detailed understanding of its relationship with other network topological properties is still limited. In fact, the task of clearly identifying the true community structure within an empirical network is complicated by a multiplicity of community detection algorithms, multiple and conflicting definitions of communities, inconsistent outcomes from different approaches, and a relatively small number of networks for which ground truth is known. Although node attributes in empirical networks (e.g., habitat type in food-webs) are sometimes used to evaluate the accuracy of community detection methods [12], these results are generally of ambiguous value as the failure to recover communities that correlates with some node attribute may simply indicate that the true features driving the network’s structure are unobserved, not that the identified communities are incorrect.

A more straightforward method of exploring the structural and functional role of a network property is to generate graphs which are random with respect to other properties except the one of interest. For example, network properties such as degree distribution, assortativity and clustering coefficient have been studied using the configuration model [13], and models for generating random graphs with tunable structural features [2,14,15]. These graphs serve to identify the network measures that assume their empirical values in a particular network due to the particular network property of interest. In this work, we propose a model for generating simple, connected random networks that have a specified degree distribution and level of community structure.

Random graphs with tunable strength of community structure can have several purposes such as: (1) serving as benchmarks to test the performance of community detection algorithms; (2) serving as null models for empirical networks to investigate the combined effect of the observed degrees and the latent community structure on the network properties; (3) serving as proxy networks for modeling network dynamics in the absence of empirical network data; and (4) allowing for the systematic study of the impact of community structure on the dynamics that may flow on a network. Among these, the use of random graphs with tunable strength of community structure to serve as benchmarks has received the most attention and several such models have been proposed [16–21]. A few studies have also looked at the role of community structure in the flow of disease through contact networks [22–25]. However, the use of modular random graphs, which can be defined as random graphs that have a higher strength of community structure than what is expected at random, is still relatively unexplored in other applications.

## Previous work

In 2002, Girvan and Newman proposed a simple toy model for generating random networks with a specific configuration and strength of community structure [3]. This model assumes a fixed number of modules each of equal size and where each node in each module has the same degree. In this way, each module is an Erdős-Rényi random graph. To produce modular structure, different but fixed probabilities are used to produce edges within or between modules. Although this toy model has been widely used to evaluate the accuracy of community detection algorithms, it has limited relevance to real-world networks, which are generally both larger and much more heterogeneous. Lancichinetti et al [16] introduced a generalization of the Girvan-Newman model that better incorporates some of these features, e.g., by including heterogeneity in both degree and community size. However, this model assumes that degrees are always distributed in a particular way (like a power law [26]), which is also unrealistic. (A similar model by Bagrow [17] generates modular networks with power law degree distribution and constant community size.)

Yan et al [23] used a preferential attachment model to grow scale-free networks comprised of communities of nodes whose degrees follow a power-law distribution. And, models for special graph types such as hierarchical networks [18], bipartite networks [21], and networks with overlapping modules [20] have also been proposed. These models also make strong assumptions about the degree or community size distributions, which may not be realistic for comparison with real biological networks. A recently proposed model [19] does generate networks with a broad range of degree distributions, modularity and community sizes, but its parameters have an unclear relationship with desired properties (such as degree distribution and modularity), making it difficult to use in practice. Thus, while these models may be sufficient for comparative evaluation of community detection algorithms, they are of limited value for understanding their performance and output when applied to real-world networks.

An alternative approach comes from probabilistic models, of which there are two popular classes. Exponential random graph models (ERGMs) have a long history of use in social network analysis, and can generate an ensemble of networks that contain certain frequencies of local graph features, including heterogeneous degrees, triangles, and 4-cycles [27]. However, many classes of ERGMs exhibit pathological behavior when parameterized with triangles or higher-order structures [28], which severely limits their utility. Stochastic block models (SBMs) are more promising, but require a large number of parameters to be chosen before a graph can be generated. In this approach, the probability of each link depends only on the community labels of its endpoints. Thus, to generate a network, we must specify the number of communities *K*, their sizes (in the form of a labeling of the vertices), and the 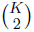 (in the undirected case) group-pair probabilities. The result is a random graph with specified community sizes, where each community is an Erdős-Rényi random graph with a specified internal density, and each pair of communities is a random bipartite graph with specified density. The degree distributions of these networks is a mixture of Poisson distributions, which can be unrealistic. A recent generalization of the SBM due to Karrer and Newman [29] allows the specification of the degree sequence, which circumvents this limitation but introduces another set of parameters to be chosen. Although the stochastic block models can in principle be used to generate synthetic networks, they are more commonly used within an inferrential framework in which community structure is recovered by estimating the various parameters directly from a network. As a result, the practical use of the SBM as a null model, either for general benchmarking of community detection algorithms or for understanding the structure of biological networks, remains largely unexplored, and we lack clear answers as to how best to sample appropriately from its large parameter space in these contexts. The SBM also does not provide a simple measure of the level of modularity in a network’s large-scale structure, which makes its structure more difficult to interpret. The SBM is a promising model for many tasks, and adapting it to the questions we study here remains an interesting avenue for future work.

## Our approach

Here, we develop and implement a simple simulation model for generating modular random graphs using only a small number of intuitive and interpretable parameters. Our model can generate graphs over a broad range of distributions of network degree and community size. The generated graphs can range from very small (*<* 10^2^) to large (*>* 10^5^) network sizes and can be composed of a variable number of communities. In Methods below, we introduce our algorithm for generating modular random graphs. In Results, we consider the performance of our algorithm and structural features of our generated graphs to show that properties such as degree assortativity, clustering, and path length remain unchanged for increasing modularity. We next demonstrate the applicability of the generated modular graphs to test the accuracy of extant community detection algorithms. The accuracy of community detection algorithms depends on several network properties such as the network mean degree and strength of community structure, which is evident in our analysis. Finally, using a few empirical biological networks, we demonstrate that our model can be used to generate corresponding null modular graphs under two different models of randomization. We conclude the paper with some thoughts about other applications and present some future directions.

## Methods

We present a model that generates undirected, simple, connected graphs with prescribed degree sequences and a specified level of community structure, while maintaining a graph structure that is otherwise as random (uncorrelated) as possible. Below, we introduce some notation and a metric for measuring community structure, followed by a description of our model and the steps of the algorithm used to generate graphs with this specified structure.

### Measure of community structure

We begin with a graph *G* = (*V, E*) that is comprised of a set of vertices or nodes *V* (*G*) = {*v*_1_, …, *v_n_*} and a set of edges *E*(*G*) = {*e*_1_, …, *e_m_*}. *G* is undirected and simple (i.e. a maximum of one edge is allowed between a pair of distinct nodes, and no “self” edges are allowed). The number of nodes and edges in *G* is *|V* (*G*)| = *n* and *|E*(*G*)| = *m*, respectively. The neighborhood of a node *v_i_* is the set of nodes *v_i_* is connected to, *N* (*v_i_*) = {*v_j_* | (*v_i_*, *v_j_*) ∈ *E*, *v_i_* ≠ *v_j_*, 1 ≤ *j* ≤ *n*}. The degree of a node *v_i_*, or the size of the neighborhood connected to *v_i_*, is denoted as *d*(*v_i_*) = *|N* (*v_i_*)|. A *degree sequence*, *D*, specifies the set of all node degrees as tuples, such that *D* = { (*v_i_*, *d*(*v_i_*)} and follows a probability distribution called the *degree distribution* with mean 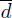.

Each community or *module C_k_* is defined as a subset of *G* that contains both nodes, *V* (*C_k_*) and edges *E*(*C_k_*), where both the endpoints of each edge in *E*(*C_k_*) are contained in *V* (*C_k_*). *K* is the number of modules in *G* and k ∈ [1, *K*]. Each node *v_i_* of *G* has a *within-degree*, *d_w_* (*v_i_*) = *|N* (*v_i_*) *∩ V* (*C_k_)*|, which is the number of *within-edges* connecting *v_i_* to other nodes of the same module *C_k;_* and a *between-degree*, *d_b_*(*v_i_*) = *|N* (*v_i_*) – *V* (*C_k_)*|, i.e. the number of *between-edges* connecting *v_i_* to nodes in different modules (here, the minus operator represents set difference). The strength of the community structure defined by a partition, {*C_k_* }, can be measured as *modularity*, *Q*, and is defined as

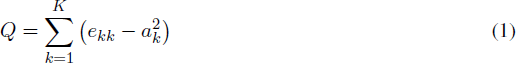

where 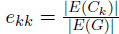 denotes the proportion of all edges that are within module *C_k,_* and 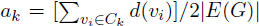 represents the fraction of all edges that touch nodes in community *C_k_*. When *Q* = 0, the density of within-community edges is equivalent to what is expected when edges are distributed at random, conditioned on the given degree sequence. Values approaching *Q* = 1, which is the maximum possible value of *Q*, indicate networks with strong community structure. Typically, values for empirical network modularity fall in the range from about 0.3 to 0.7 [30]. However, in theory Good et al. [31] show that maximum *Q* values depend on the network size and number of modules.

In order to generate a graph with a specified strength of community structure, *Q*, equation (1) represents our first constraint, which we rewrite below in terms of the expected value of *Q*, (full derivation in Additional file 1):

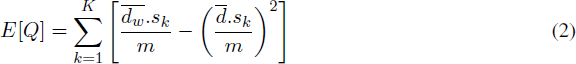

where 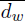 and 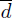 are the average within-degree and average degree, respectively, and *s_k_*= *|V* (*C_k_)*| is the module size for module *k*. Thus, equation (2) allows us to specify 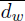 in terms of *Q*, 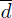, *m* and *sk_,_* assuming that the module-specific average degree and average within-degree are equal to 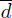 and 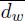, respectively. When 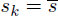 for all *k*, *E*[*Q*] reduces to 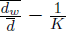.

We note that as the average within-degree 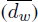 approaches the average degree 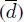, the graph, *G* becomes increasingly modular. Hence, the maximum modularity for *G* with *K* modules can be estimated as:

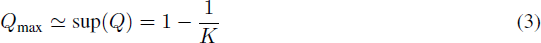

### Algorithm

We present a model and an algorithm that generates undirected, unweighted, simple and connected modular random graphs. The model is specified by a network size (*n*), degree distribution (*p_d_*), an expected modularity (*E*[*Q*]), the number of modules (*K*), and the module size distribution (*P* (*s*)), with mean 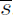. (We note a degree sequence, *d*(*v_i_*), may be specified instead of a degree distribution, *p_d_*). The algorithm proceeds in four steps:

1. Assign the *n* network nodes to *K* modules based on the size distribution *P* (*s*).
2. Assign degrees, *d*(*v_i_*), to each node *v_i_* based on *p_d_* and 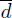. We next assign within-degrees, *d_w_* (*v_i_*), to each node *v_i_* by assuming that the within-degrees follow the same distribution as *p_d_* with mean 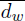, which is estimated based on equation (2) above (Figure 1a).
3. Connect between-edges based on a modified Havel-Hakimi model and randomize them. (Figure 1b)
4. Connect within-edges based on the Havel-Hakimi model and randomize them. (Figure 1c and 1d)

The generated graph then has a degree distribution that follows *p_d_* with mean 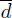, *K* modules with sizes distributed as *P* (*s*), and a modularity *Q ≈ E*[*Q*]. We set an arbitrary tolerance of ∈ = 0.01, such that the achieved modularity is *Q* = *E*[*Q*] *±* ∈. The graph is also as random as possible given the constraints of the degree and community structure, contains no self loops (edges connecting a node to itself), multi-edges (multiple edges between a pair of nodes), isolate nodes (nodes with no edges), or disconnected components. Below, we elaborate on each of the steps of this algorithm.

**Figure 1.**
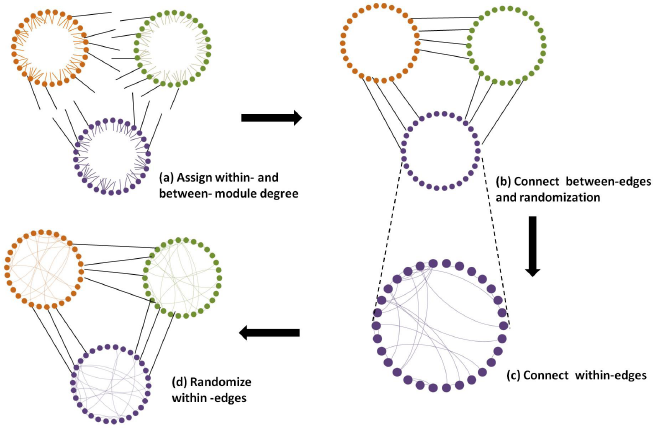
Schematic representation of the steps of our algorithm. **(a)** The algorithm assigns a within-degree and between-degree to each node, which are represented here as half-within-edges and half-between-edges respectively. **(b)** The half-between-edges are then connected using a modified version of the Havel-Hakimi algorithm, and to remove degree correlations, the between-edges are randomized. Finally, the half-within-edges are connected using the standard Havel-Hakimi algorithm for each module and **(d)** the within-edges are randomized to remove degree correlations.

#### Assigning nodes to modules

We sample module sizes, *s_k_*, for each of the *K* modules from the specified module size distribution, *P* (*s*) so that 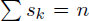. The *n* nodes are then arbitrarily (without loss of generality) assigned to each module to satisfy the sampled module size sequence.

#### Assigning degrees

Based on the degree distribution specified, a degree sequence is sampled from the distribution to generate a degree, *d*(*v_i_*), for each node *v_i_* (unless a degree sequence is already specified in the input). To ensure that the degree sequence attains the expected mean of the distribution (within a specified threshold) and is *realizable*, we verify the Handshake Theorem (the requirement that the sum of the degrees be even) and the Erdős-Gallai criterion (which requires that for each subset of the *k* highest degree nodes, the degrees of these nodes can be “absorbed” within the subset and the remaining degrees) [32], and that no node is assigned a degree of zero.

Unless a within-degree sequence is specified, we assume that the within-degree distribution follows the class of the degree distribution specified, *p_d_*, with mean 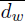 based on equation (2) (i.e. a generated network with a Poisson degree distribution of mean 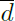 also has a Poisson within-degree distribution with mean 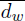). This assumption is considered reasonable as it holds true for several of the empirical networks we analyze (shown in Figure S1 in Additional file 1). However, our model can be extended for arbitrary within-degree distributions (or sequences) (see Table S1 in Additional file 1), although the space of feasible within-degree distributions given a degree distribution is restricted. Next, we sample a within-degree sequence, *d_w_* (*v_i_*), from this within-degree distribution. Using rejection sampling, we ensure that the within-degree sequence attains the expected overall mean, 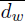 within a tolerance 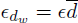 (with details in the Additional file 1), and satisfies the following conditions:

*•* Condition 1: *d*(*v_i_*) ≥ *d_w_* (*v_i_*) for all *v_i_*. To ensure this, we sort the degree sequence and within-degree sequence, independently. If *d*(*v_i_*) < *d_w_* (*v_i_*) for any *v_i_* in the ordered lists, the condition is not satisfied. In Figure S2 of Additional file 1, we discuss the rejection rates for the rejection sampling of both the degree and within-degree sequence.

*•* Condition 2: a realizable within-degree sequence for each module, *C_k_*, as defined by the Handshake Theorem and the Erdos-Gallai criterion.

In addition, to ensure that each module approximately achieves the overall mean within-degree, 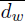, we specify the following constraint: max[{*d_w_*(*v_i_*)}*v_i_*∈*G*] *≤* min[*s_k_*]. If the sampled module sizes do not satisfy this criteria, the module sizes are re-sampled or an error is generated.

The between-degree sequence is generated by specifying *d_b_*(*v_i_*) = *d*(*v_i_*) − *d_w_* (*v_i_*) for each node *v_i_*. To test if the between-degree sequence is realizable, we impose a criterion developed by Chungphaisan [33] (reviewed by Ivanyi [34]) for realizable degree sequences in multigraphs. To do so, we imagine a coarse graph, *H*, where the modules of *G* are the nodes of *H* (i.e. *V* (*H*) = {*C*_1_, *C*_2_,… *C_K_* }), and the between-edges that connect modules of *G* are the edges of *H*. We note that *H* is a multigraph, because *G* allows multiple between-edges of *G* to connect each pair of modules. In this case, the degree sequence of *H* is 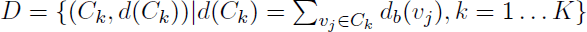.

The Chungphaisan criterion then specifies that the multigraph degree sequence {*d*(*C_k_*)} on *H* is realizable if the following conditions are satisfied:

- Condition 1: the Handshake theorem is satisfied for 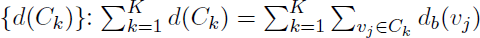 is even
- Condition 2: 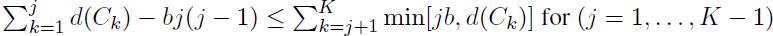.

Here, *b* is defined as the maximum number of edges allowed between a pair of nodes in *H*; in our case, *b* = max[{*d_b_*(*v_i_*)}], the maximum between-degree of any node *v_i_* ∈ *G*.

We also generate graphs with *Q* = 0 by assuming the network is composed of a single module with no between-edges. Thus, *d_w_* (*v_i_*) = *d*(*v_i_*) and *d_b_*(*v_i_*) = 0 for all *v_i_* ∈ *G*.

#### Connecting edges

Based on the within-degree sequence and between-degree sequence specified above, edges are connected in two steps (Figure 1). Nodes that belong to different modules are connected based on their between-degree to form between-edges (Figure 1b) and nodes that belong to the same module are connected according to their within-degree to form within-edges (Figure 1c and 1d).

We connect between-edges using a modified version of the Havel-Hakimi algorithm. The Havel-Hakimi algorithm [35,36] constructs graphs by sorting nodes according to their degree and successively connecting nodes of highest degree with each other. After each step of connecting the highest degree node, the degree list is resorted and the process continues until all the edges on the graph are connected. Here, we modify this to construct between-edges by sorting nodes by highest between-degree, in order of highest total between-degree for the module to which they belong, and successively connecting the node at the top of the list randomly with other nodes. Connections are only made between nodes if they are not previously connected, belong to different modules, and do not both have within-degree of zero (to avoid disconnected components). After each step the between-degree list is resorted, and the process continues until all between-edges are connected. After all between-edges have been connected, the connections are randomized using a well-known method of rewiring through double-edge swaps [37]. Specifically, two randomly chosen between-edges (*u, v*) and (*x, y*) are removed, and replaced by two new edges (*u, x*) and (*v, y*), as long as *u* and *x*, and *v* and *y* belong to different modules, respectively. The swaps are constrained to avoid the formation of self loops and multi-edges. This process is repeated a large number of times to randomize edges.

We then connect within-edges using the standard Havel-Hakimi algorithm, applied to each module independently. Specifically, within-edges of a module are connected by sorting nodes of the module according to their within-degree and successively connecting nodes of highest within-degree with each other. Connections are only made between nodes if they are not previously connected, and do not both have a between-degree of zero (to avoid disconnected components). After each step the within-degree list is resorted and the process continues until all the within-edges of the module are connected. The connections are then randomized by rewiring through double-edge swaps [37]. We do not specify that each module be connected (only that the full graph is connected). However, if this is required, Taylor’s algorithm can be used to rewire pairs of edges until the module is connected [38]. Specifically, the algorithm selects two random edges (*u, v*) and (*x, y*) that belong to two different disconnected components of the module. As long as (*u, x*) and (*v, y*) are not existing edges, the (*u, v*) and (*x, y*) edges are removed and (*u, x*) and (*v, y*) are added. Taylor’s theorem proves that following such operation any disconnected module can be converted to a connected module with the same degree sequence.

## Results and discussion

Using our simulation algorithm, we were able to generate modular random graphs of variable network size, number of communities, degree distribution, and community size distribution. In Figure 2, we show sample networks of varying levels of modularity, *Q* = 0.1, 0.3, 0.6. We note that a network with three modules can approach a maximum modularity value of 2/3 (from equation 3), and thus *Q* = 0.6 is a relatively high modularity for this particular network type. In the sections that follow, we consider the algorithm performance, as well as structural properties of the generated graphs. We then highlight two applications of our model: 1) to generate benchmark graphs for validation of community detection algorithms and 2) to generate null graphs for the analysis of empirical networks. Community detection algorithms assist in identifying community structure in empirical networks. Our model is able to generate modular networks *de novo* to test these algorithms. Once community structure has been identified in an empirical network with a community detection algorithm, the number of communities and the modularity level (*Q*) (and, if desired, the community size distribution and within-degree sequence) can be used as input to our model to generate graphs that can act as random controls to test hypotheses about the empirical system.

**Figure 2.**
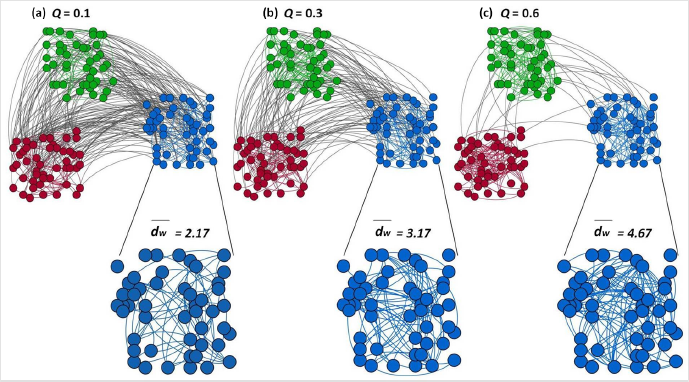
Modular random graphs with *n* = 150, *m* = 375, *K* = 3, *P* (*s* = 50) = 1 and *pk* is power law with modularity values of : a) *Q* = 0.1; b) *Q* = 0.3; and c) *Q* = 0.6. As the modularity increases, the ratio of the total number of edges within modules to the number of edges in the network increases (i.e. 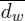 increases), while the remaining parameter values (degree distribution, network mean degree, number of modules) are held constant.

### Performance & properties of generated graphs

#### Performance

Our model generates graphs that closely match the expected modularity and degree distribution. The deviation of the observed modularity is less than 0.01 from the expected value, given the specified partition. The modular random graphs with Poisson degree distribution generated by our model are similar to the ones described by Girvan and Newman [3] with linking (*p_in_*) and cross-linking probability (*p_out_*) equal to 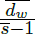 and 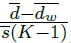 respectively. However, our model overcomes several limitations of the model proposed by Girvan and Newman [3] and others [16,17] by considering heterogeneity in total degree, within-module degree distribution, and module sizes. Unlike many of the existing models [18–21], our model can generate modular random graphs with arbitrary degree distributions, including those obtained from empirical networks. Though we discuss modular random graphs with positive *Q* values, our model can also generate disassortative modular random graphs (see Figure S3 in Additional file 1). In this case, nodes tend to connect to nodes in other modules and thus the density of edge connections within a module is less than what is expected at random. Additionally, we also compare our model to graphs generated based on a degree-corrected stochastic block model (SBM). The details of the parameterization of the SBM and the results are shown in Figure S4 in Additional file 1).

#### Structural properties

There are several other topological properties (besides degree distribution and community structure) that can influence network function and dynamics. The most significant of these properties are degree assortativity (the correlation between a node’s degree and its neighbor’s degrees), clustering coefficient (the propensity of a node’s neighborhood to also have edges among them) and average path length (the typical number of edges between pairs of nodes in the graph). We have developed this model to generate graphs with specified degree distribution and modularity, while minimizing structural byproducts. Thus, it is important to confirm that we have reached this goal with the generative model above.

To evaluate the status of other structural properties due to the generative model, we specify graphs of *n* = 2000 following Poisson (λ*^k^ e^-λ^/k*!), geometric (*p*(1 − *p*)^*k*−1^), and power-law 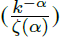 degree distributions with 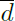 = 10. We chose these particular types of degree distributions as they have widely studied in the context of biological networks [39–41]. Each network has *K* = 10 modules and a module size distribution *P* (*s* = 200) = 1. We generate modular random graphs with these specifications and modularity values that range from *Q* = 0 to *Q* = 0.8, in steps of 0.1. For each level of modularity, we generated 50 such modular random graphs and calculated the degree assortativity (*r*), clustering coefficient (*C*), and average path length (*L*) for each network, which is illustrated in Figure 3. In networks with random community structure (*Q* = 0), that is random graphs with specified degree distributions (such as those that would be generated by the configuration model [13]), the value of *r, C,* and *L* are what are expected at random. In Figure 3, we show that for increasing values of modularity, degree assortativity, clustering coefficient, and average path length remain relatively constant for all three network types (i.e. Poisson, geometric and power-law). At the highest levels of modularity, edge connections are constrained, particularly for the heavy-tailed geometric and power-law degree distributions, leading to an increase in clustering coefficient. Correlations between high clustering coefficient and high modularity have also been observed before [2]. The average path length for all network types also increases at the highest levels of modularity, likely reflecting the lack of many paths between modules, requiring additional steps to reach nodes in different modules. Thus, our model is able to increase levels of modularity in random graphs without altering other topological properties significantly.

**Figure 3.**
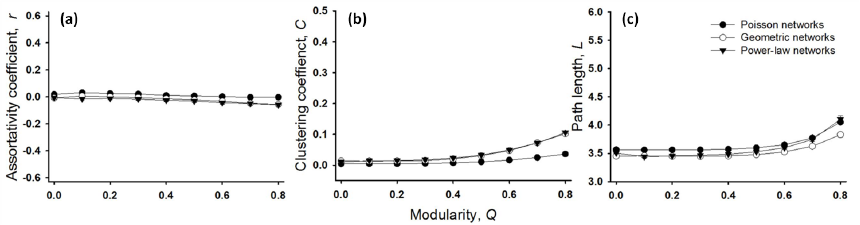
Values of (a) Assortativity, *r*, (b) clustering coefficient,*C*, and (c) path length, *L* in modular random graphs do not vary significantly with increasing modularity (*Q*). Each graph has *n* = 2000 nodes, a mean degree 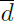 = 10 and *K* = 10 modules with *P* (*s* = 200) = 1. The data points represent the average value of 50 random graphs. Standard deviations are plotted as error bars.

Biological networks show remarkable variation in network size, connectivity and community size distribution, with some of them having particularly small network size, high degree, and small module sizes (e.g. food-web networks). We therefore tested the performance of our generated networks under deviations in the network specifications of size, mean degree and module size distribution (results presented in Additional file 1: Figure S5, S6 and S7). We find that the structural properties of our generated modular random graphs remain constant, except for two constraining conditions: a) high average degree 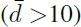 and b) low average module size 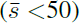. At these parameter extremes, the modular random graphs become degree disassortative and have increased clustering coefficient. A similar observation of network degree disassortativity has been made in hierarchically modular networks [42]. In these two scenarios, the highest value of within-degree (*d_w_* (*v_i_*)) that a node can attain is constrained by the community size, which reduces the number of possible high within-degree nodes. As a consequence high within-degree nodes must connect to low within-degree nodes more than expected, resulting in a degree disassortative network. In these two cases, modules also become more dense and thus create more triangles resulting in a gradual increase in clustering. Path length, on the other hand, is not affected by these conditions and shows a consistent dependence on network size and mean degree, which is well known [43,44].

### Application: Benchmark graphs for community-detection algorithms

Detecting communities in empirical networks has been an area of intensive research in the past decade [45] since Girvan and Newman’s seminal paper on community detection [3]. Extant techniques such as modularity maximization, hierarchical clustering, the clique-based method, the spin glass method etc. aim at achieving high levels of accuracy in detecting the correct partition (for a detailed review see [45]), but have their own set of strengths and weaknesses. Choosing the best algorithm can be a difficult task especially as algorithms often use distinct definitions of communities and perform well within that description. Thus, it is exceedingly important to test community-detection algorithms against a suitable benchmark. We propose our modular random graphs as benchmark graphs for the validation of existing and new algorithms of community detection.

To illustrate this use, we test the performance of six popular community detection algorithms: the Louvain method [46], fast modularity method [47], the spin-glass based method [48], the infoMAP method [49], label propagation [50] and the random-walk based method [51] using our modular random graphs as benchmarks. Specifically, we generate a modular random graph for each level of modularity and used these community detection algorithms to detect their community structure. We also test the performance of the algorithms on random graphs of specified degree distribution, with no modularity (i.e. *Q* = 0). Figure 4 summarizes the performance of these algorithms, as measured by the estimated *Q*, for modular random graphs with three different degree distributions (Poisson, geometric and power-law). We also investigated the robustness of these algorithms on replicate modular random graphs at each modularity level, with the results presented in (Additional file 1: Figures S8, S9 and S10).

**Figure 4.**
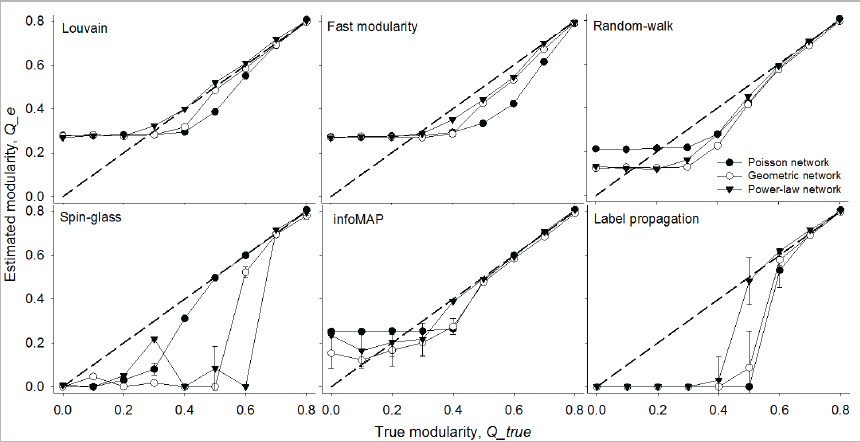
Performance of the Louvain method [46], fast modularity method [47], the spin-glass based method [48], the infoMAP method [49], label propagation [50] and the random-walk based method [51] in networks with mean degree 10. Fill circles, open circles and triangles represent networks with Poisson, geometric and power-law degree distributions, respectively. Each data point represents the average over ten modular random graphs. Error bars represent standard deviations. The solid line is the reference line where estimated modularity is equal to the input modularity.

The Louvain, fast modularity algorithm, random-walk and infoMAP algorithm overestimate the modularity for networks with weak community structure, and underestimate the modularity for networks of moderate community structure across all three network types (Figure 4). Spinglass and label-propagation consistently underestimate the modularity of both weak and moderate community structure. All the algorithms are fairly accurate at the highest strengths of community structure across the various network types. The accuracy at a particular level of modularity and degree-distribution, however, varies for different algorithms. For instance, the performance of spin-glass algorithm is better for Poisson modular random graphs at modularity values of 0.5-0.6, whereas the Louvain and label-propagation algorithm out-perform on geometric random modular graphs at these modularity values.

In addition to comparing the estimated values of modularity to the known values in the modular random graphs, we can compare the similarity in the partitions detected by the algorithms to the true partitions. For this comparison, we use the Jaccard similarity (*J*), which measures the similarity between two partitions based on the proportion of the union of the partitions that is made up by the intersection of the partitions [52]; as well as the *Variation of Information (VI*), which measures the distance between two partitions based on the amount of information lost when going from one partition to another [53]. These results are presented in the (Additional file 1: Figure S11). As reflected in the results above, we find that partitioning is inaccurate when the true community structure is weak but improves as the *Q_true_* value increases. These observations have also been noted before by Lacichinetti and Fortunato [54].

### Application: Null analysis of empirical networks

It is crucial to have random controls in the study of biological systems. Our algorithm can be used to generate null models and applied to the detection of structure in empirical biological networks. These null networks can be used to test hypotheses regarding the role of modularity and other topological features of the empirical networks. To do so, one would first determine the number of communities and modularity level (*Q*) of the sampled network using an appropriate community detection algorithm (the previous section describes the use of random modular graphs to validate existing algorithms of community detection). Our algorithm can then be used to generate an ensemble of networks that match the empirical degree structure and community structure, and then compare the structural, functional, or dynamical properties of the empirical network to those of the generated modular random graphs. Because our model generates graphs without any structural byproducts (as illustrated in a previous section), this is an appropriate model for generation of null models. We note that our algorithm does not necessarily require knowledge of the complete empirical network, but rather only estimates of the degree structure and community structure. The literature on algorithms for inference of network structure from a sample is growing, and currently includes work on inference of missing nodes, edges and even community structure [55–57].

We demonstrate this application using four classes of biological networks, namely: a) a food-web, representing the trophic interactions at Little Rock Lake in Wisconsin with a network size of 183 and average degree = 26.8 [58]; b) a protein-protein interaction network in *Saccharomyces cerevisiae* (a yeast) of size= 4713 and average degree = 6.3 [59]; c) a metabolic interaction network of *Caenorhabditis elegans* of size = 453 and average degree 9.0 [60]; and d) a network of social interactions in a community of dolphins living off Doubtful Sound, New Zealand of size = 62 and average degree = 5.1 [61]. Visualizations of the dolphin social interaction network and the food-web trophic interaction network and its modular random counterpart are shown in Figure 5.

**Figure 5.**
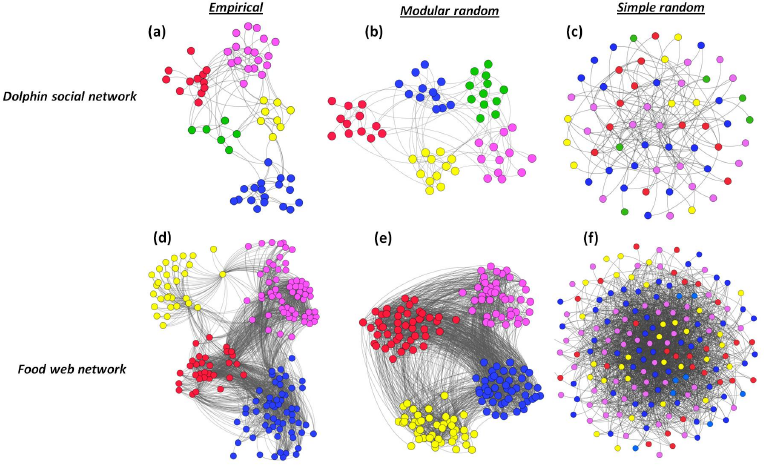
Visualization of empirical and random graphs of social interaction of dolphins and food-web trophic interactions at the Little Rock Lake in Wisconsin. Figure **(a)** is the empirical network of Dolphin social network, **(b)** its modular random graph, and **(c)** its random graph counterpart with matched degree distribution (*Q* = 0*b*). Figure **(d)** is the empirical network for the food-web trophic interaction at Little Rock Lake in Wisconsin, **(e)** is its modular random graph and **(f)** its random graph counterpart with matched degree distribution. Modular random graphs have generated to match the overall degree distribution, network mean degree, the level of modularity and the number of modules of the empirical graphs. Random graphs with matched degree distribution are based on the configuration model.

For each of these four empirical networks, we generate modular random graphs (Figure 6, light gray bars) with three parameters estimated from the empirical networks: (a) the degree sequence, *pk* (b) the modularity, *Q* and (c) the average community size, 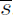. We note that as our goal is to construct null models, we assume that communities are of equal size, i.e. 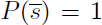, and that the within-degree distribution matches the degree distribution fitted from the specified degree sequence (with estimated mean, 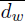). (A second class of null models can be constructed with *P* (*s*) and the within-degree sequences estimated from the empirical networks, and we do this in Table S2 of Additional file 1). Specifically, we generate 25 such random graphs and measure structural properties of the generated graphs including clustering coefficient (*C*), average path length (*L*), degree assortativity (*r*).

**Figure 6.**
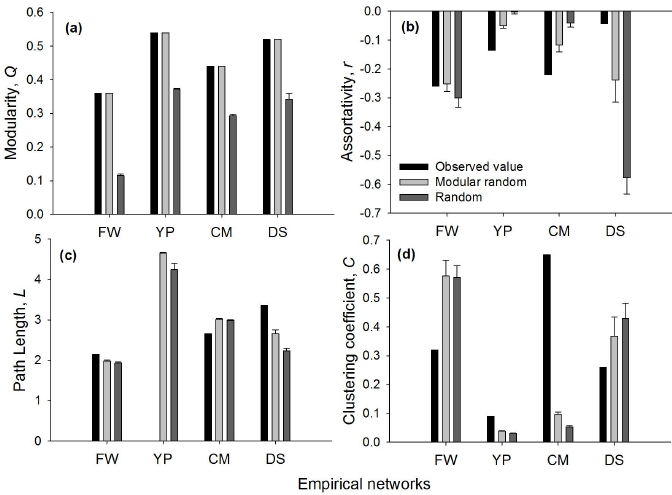
Comparisons of empirical networks, modular random graphs and random graphs with matched degree distribution (based on the configuration model). The figure summarizes network statistics of the empirical network as well as the ensemble mean of two types of random graphs in terms of (a) Modularity, *Q;* (b) Assortativity, *r*; (c) Path length, *L* and (d) Clustering coefficient, *C*. The path length value for the empirical Yeast-Protein interaction network is missing as the network contains disconnected components. Error bars denote standard deviation from the ensemble mean of the generated random graphs. Errors bars for modular random graphs in Figure 6(a) have been omitted as the value of modularity (*Q*) match the empirical networks perfectly. FW = Little Rock food web, YP = Yeast protein interaction network, CM = *C.elegans* metabolic network and DS = Dolphin social network.

We also generate random graphs based on the configuration model that have the same degree distribution and average network degree as the empirical network but are random with respect to other network properties for each of the four empirical networks (Figure 6, dark gray bars). Our modular random graph model identifies which network measures assume their empirical values in a particular network because of (i) the observed degrees and (ii) the latent community structure. The configuration model, on the other hand, only specifies (i) and not (ii) [13]. Comparison to these configuration model networks thus helps us highlight the utility of our model to identify which empirical patterns in a network are deserving of further investigation. Figure 6 shows the value of each of these properties for the empirical networks as well as the ensemble mean of modular random and random graphs with matched degree distribution.

From Figure 6 it is evident that none of the empirical biological networks have network structure identical to their null counterparts. This suggests that the structure of each of these biological systems is governed by more than what is specified by the degree distribution and community structure. However, the observed network properties of empirical networks are closer to the ensemble means of the modular random graphs, which indicates that modularity is an essential structural component of real biological networks and that it plays an important role in influencing other structural properties of the network. For instance, compartmentalization induced by modularity promotes species persistence and system robustness by containing localized perturbation [11,62,63], which might favor their selection during the course of evolution. Our results show that the empirical networks tested have a much higher modularity than the simple random graphs (Figure 6a) and therefore provide evidence for this selection. Out of the three network properties that we tested apart from modularity, we found clustering coefficient of the generated random graphs to be significantly different from each of the empirical counterparts. This may point to a functional role for “triangles” in these biological networks, significantly above or below what is prescribed by the degree and community structure.

#### Little Rock Lake food web interactions (FW)

Among the four empirical networks that we tested, the properties of the ensemble mean of null models such as assortativity and path length closely match most of the observed properties of Little Rock food web. The observed clustering coefficient of food web is strikingly lower than either of the random graphs which confirm the observations of low clustering in food web made by earlier studies (Figure 6d). The observed path length of this food web is short (Figure 6c) and only slightly longer than the path lengths of random graphs, which has also been noted before [64–66]. We note that for this food web, the structural properties of the random graphs with matched degree distribution are quite similar to those of modular random graph counterparts, suggesting that the degree distribution, particularly the high density of edges in the network governs most of the other topological characteristics of this network. Modularity, on the other hand, seems to play a minor role in dictating the structural properties of this network.

#### Yeast protein-protein interaction network (YP)

The empirical yeast protein network is more disassortative than the ensemble mean of null modular graphs (Figure 6b). Disassortative interactions in protein-protein interaction networks are known to reduce interferences between functional modules and thus increase the overall robustness of the network to deleterious perturbations [6], while also allowing for functions to be performed concurrently [67]. The results therefore suggest that disassortative interactions may be selected for in the evolution of biological networks. From Figure 6(d) it is also evident that the yeast protein network has a higher value of clustering coefficient than the expected value predicted by the modular random graphs. A high value of clustering coefficient indicates that there are several alternate interaction paths between two proteins, making the system more robust to perturbation [68].

#### *C. elegans* metabolic interaction network (CM)

The *C.elegans* metabolic network demonstrates a shorter path length but higher clustering coefficient than both modular and random graphs with matched degree distribution (Figure 6c and 6d). A high clustering coefficient and short path length suggests that the graph has small-world properties, which has been observed in other metabolic networks as well [69]. A highly disassortative degree structure is also well known in metabolic networks, although the mechanism leading to this property is unclear (see review by [39]). As the predicted value of disassortativity of the modular random graphs is closer to the observed value, our results suggest that the strong community structure of the metabolic networks could be one of the factors contributing to high degree disassortativity. (As discussed earlier, community structure leads to significant degree correlations in small networks with long-tailed degree distributions; see Figure S5 in Additional file 1 for an example).

#### Social interaction network of dolphins network at Doubtful Sound, New Zealand (DS)

The empirical social interaction network of dolphins that we investigated demonstrated a negative assortativity (or disassortativity) similar to other real biological networks (Figure 6b). Interestingly, the assortativity value of both null modular and random graphs with matched degree distribution counterparts of the dolphin network is *lower*than the observed value, which suggests that the network is more assortative than expected. Degree assortativity has also been observed in other animal [70] and human [4] social interaction networks. This result is quite intuitive for a social network and is also referred to as homophily: more gregarious individuals tend to interact with other gregarious individuals while introverted individuals prefer to associate with other introverts [14]. The empirical dolphin network also demonstrated a lower value of clustering coefficient than the expected values of either null model. Low clustering coupled with high degree assortativity indicates that dolphin populations may be more susceptible to the propagation of infection or information, as transmission may occur rapidly through the entire network with such properties [70,71].

## Conclusions

In summary, the model that we propose in this study generates modular random graphs over a broad range of degree distribution and modularity values, as well as module size distributions. We highlight that our model is specifically designed to generate networks which have modularity evenly divided across its modules, modulo the impact of module size. This means that we are mitigating the resolution limit effect and indeed generating networks with the maximum modularity partition. We also confirm that structural properties of our generated modular graphs such as assortativity, clustering and path length remain unperturbed for a broad range of parameter values. This important feature allows these graphs to act as benchmark and control graphs to explicitly test hypotheses regarding the function and evolution of modularity in biological systems. Of the approaches available, our method provides flexibility and has been explored the most fully for these applications.

Compartmentalization of biological networks has been an area of great interest to biologists. What we refer to as community structure in this work is any segregation of a biological system into smaller subunits inter-connected by only a few connections. It has been suggested that modularity in a system promotes system robustness and enhances species persistence by containing localized perturbations [11, 63]. Metabolic networks of organisms living in a variable environment have indeed been found to be more modular [62]. Maintaining and selecting for modularity in biological networks, however, comes at a great cost of reducing system complexity [72], longer developmental time and cost of complete module replacement in case of failure [73]. It is therefore unclear why modularity would be strongly selected for as a structural feature of biological systems. There is also a lack of evidence to prove that the functional localization of sub-goals overlaps with the structural segregation of the network into community structure. Our work provides a tool for the systematic study of network structure (through benchmark graphs) and of the impact of connectivity and compartmentalization on system function and dynamics (through control graphs).

The detection of community structure plays a crucial role in our topological understanding of complex networks. Currently the performance of community detection methods is usually evaluated based on ground-truth from real networks. However, determining reference communities in real networks is often a difficult task. Also, ground truth data on empirical network partitions do not necessarily identify system features based on network topology and thus may create a bias when analyzing community structure. A more convenient technique of evaluating community detection method is to use artificial random graphs, but has been limited as most of the models fail to incorporate degree heterogeneity of real networks. By providing a systematic method to generate benchmark graphs, our model can aid in the development of more robust community detection algorithms, and therefore improve our topological understanding of empirical networks.

A step beyond identifying the topological presence of network communities is the understanding of its evolution as well as the functional and dynamical role of community structure. We believe this process can be facilitated by using an appropriate class of control or null graphs. As a model for generating null networks, our method joins a suite of random graph models, each contributing to a hierarchy of null models. The simplest model for generating random graphs (based on only a single parameter) is the Erdős-Rényi random graph model, which produces graphs that are completely defined by their average degree and are random in all other respects. A slightly more complex and general model is one that generates graphs with a specified degree distribution (or degree sequence) but are random in all other respects [13,74,75]. These models can be extended to sequentially include additional independent structural constraints, such as degree distribution and clustering coefficient [2], or degree structure and community structure, as we have demonstrated here. A further extension to this work will be designing models that generate random graphs with multiple structural constraints. For example, our model can be combined with the one proposed by [2] to generate random graphs with specified degree distribution as well as tunable strength of modularity and clustering coefficient.

## Availability and requirements

**Project name:** Modular random graph generator **Project home page:** http://github.com/bansallab **Operating system(s):** Platform independent **Programming language:** Python 2.7

**Other requirements:** Networkx Python package

**License:** BSD-style

Any restrictions to use by non-academics: **None**

## Author’s contributions

PS and SB contributed to algorithm design and implementation. PS, LOS, AC and SB contributed to manuscript writing. All authors read and approved the final manuscript.

## Acknowledgements

This work was supported by NSF award DEB-1216054.

## Additional file

**Additional_file_1 as PDF**

**Additional file 1: Supplementary analysis.** Additional analysis of algorithm with figures

